# Nucleotide Dynamics During Flossing of Polycation-DNA-Polycation through a Nanopore using Molecular Dynamics

**DOI:** 10.1101/2021.06.21.449276

**Authors:** Jyoti P. Mahalik, Murugappan Muthukumar

## Abstract

The full potential of solid state nanopores is yet to be realized for genome sequencing. Due to its robustness it can handle strong voltage amplitude and frequency. The effect of strong alternating voltage on the dynamics of nucleotides during translocation has been explored. We proposed a setup consisting of single stranded DNA covalently linked with symmetric polycations at both ends fashioned after the proposal of Kasianowicz. ^1^ Such a setup allows for repeated back and forth motion of the DNA along the nanopore (1.45 nm diameter and 1.53 nm thick) by simply switching the voltage polarity if the polycation tail is sufficiently long (≥ 10) and the applied voltage is below 0.72 volts, but the average residence time of the nucleotides are too small to be of any practical use (6-30 *ns*). When alternating voltage of higher frequency is applied, it enhances the average residence time of the nucleotides by an order of magnitude to ∼ 0.1 *µs* relative to direct voltage but the individual trajectories are too stochastic. Since, we are able to collect repeated read on the dynamics of individual nucleotides, we obtained the most probable time of appearance of a nucleotide within the nanopore. With such construct we were able to get almost linear dependence of most probable time versus nucleotide index, after gaussian fitting.

## Introduction

Ionic current based nanopore technology for DNA sequencing has advanced a lot in the past few decades: from research laboratories to commercial applications.^2–15^ In this technology indirect and label free analysis of DNA is done by reading the sequence of a single stranded (ss) DNA from an ionic current trace. When a ssDNA molecule translocates through a sufficiently narrow nanopore as a single file under an applied voltage, the detected ionic current is reduced from its previous level (no ssDNA inside nanopore), providing molecular signature of the translocating species. A suitable nanopore for ssDNA sequencing applications must have a sufficiently small diameter that can resolve small differences in the molecular structure of the four nucleotides through their ionic current trace. Moreover, the nanopore needs to be sufficiently thin (sub nanometer) in order to be able to resolve few nucleotides at a time. Additional requirements are not directly related to the nanopore but to the conducive environment for detection. The ssDNA must move at a deterministic rate in a particular direction while translocating through the narrowest region of the nanopore, and it must reside inside the nanopore for sufficiently long time for good signal to noise ratio (SNR). MspA protein pore has the appropriate dimension (1.2 nm diameter and 0.6 nm thickness^4^) so also many solid state nanopores such as MoS_2_ (1.4 nm diameter and 0.7 nm thickness^4^), SiN_*x*_ (1.4-2.1 nm diameter and 0.5-8.9 nm thickness^16^), and HfO_2_ (1.4 nm diameter and 2-7 nm thickness^17^). The solid state nanopores have much better SNR over protein nanopores, attributed to their capability to handle high applied voltage amplitude and frequency,^4^ yet solid state nanopores are out of reach from commercial applications. Successful commercialization of MspA protein pore for ssDNA sequencing happened due to its successful coupling with biological enzymes, which ratchets the DNA at a steady rate through MspA. This coupling not only slows down the translocation speed by orders of magnitude but also qualitatively changes the nature of motion from a stochastic state to a steady deterministic state, increasing SNR by two orders of magnitude compared to a free ssDNA translocation process.^4^ Successful attempts have been made to slow down the DNA translocation through solid state nanopores, often by orders of magnitude. Some of the methods for slowing down DNA traslocation are by using ionic liquid medium,^18^ placing tight gels on the trans side of the nanopore,^19^ putting external pressure opposing DNA motion,^20^ by mechanical trapping, ^21^ chemical modification of nanopore surface, ^22^ and nanopore charging.^23^ However, the nature of DNA translocation could not be transitioned from stochastic to deterministic, as evident from their reported wide translocation time distributions. The undesired consequence of faster and stochastic translocation on genome sequencing has been described in Ref.^24,25^

The other possible option proposed for slowing down DNA translocation through nanopores and simultaneously making its motion deterministic is by using high frequency voltage (» MHz).^4^ All atomistic molecular dynamics (MD) simulations have been used to investigate the dynamics of nucleotides of a ssDNA translocating under alternating voltage at high frequency.^26–28^ They reported oscillatory motion of nucleotides in sync with the frequency of the applied voltage. Although, the amplitude of the applied voltage in these studies were reasonable (of the order of 100 mV), the timescale of simulation were too small (∼ 1 ns^26^ and *<* 100 ns^27,28^). They clearly demonstrated that the nucleotides can be deliberately trapped inside a nanopore by using very high frequency voltage but in order to investigate whether nucleotides of a DNA molecule can be ratcheted forward at a steady rate, large time scale simulations are required. Coarse grain (CG) DNA model may be the key to ascertain the practicality of such strategies for ratcheting DNA in solid state nanopores.

Another interesting aspect of the DNA sequencing worth exploring is the capability to repeat the sequencing experiment back and forth. When a ssDNA embedded in a narrow nanopore is tethered at both ends by proteins, or DNA hairpins or segment of a base-paired DNA, it forms a dumbbell kind of structure. As suggested by Kasianowicz^1^ this dumbbell molecule can be flossed (moved back and forth through a nanopore) for increasing SNR characteristic, and thereby increasing the chance of reliably sequencing long strands of DNA with a nanopore. Such idea have been applied recently in a slightly modified form by translocating DNA through a double nanopore system. ^29,30^ In this double nanopore setup, the nanopores are in parallel and are located in close vicinity. A 48.5 kbp double stranded *λ*− phage DNA was tagged by fixed number of proteins at regular intervals and it was flossed between two nanopores by applying different bias across the two nanopores.^30^ The direction of motion of the DNA was tuned through the applied voltage; the DNA is pulled in the pore with relatively higher bias and the location of protein tags were read from the deeper ionic current levels relative to bare DNA current levels. They demonstrated that by averaging over a large number of data, the relative positions of the protein tags can be reliably determined. Molecular dynamics (MD) simulation on a coarse-grained (CG) model of a ssDNA was also done for a similar double nanopore setup.^29^ They demonstrated the practicality of capturing the DNA in the second nanopore by using hourglass shaped nanopore chambers in parallel. They also showed that by averaging over many runs, the reliability of the DNA sequence prediction increased. Along those lines, using alternating external force, stochastic resonance has been reported during the translocation of a polyelectrolyte through a nanopore.^31^ Using Fokker-Planck formalism, Mondal and Muthukumar^31^ derived an analytical expression for determining the signal to noise ratio as a function of the system parameters. For example, knowing the pore length, polymer length, and the pore-polymer interaction strength, the frequency and magnitude of the external force to detect highest signal to noise ratio can be predicted. Such an analytical formalism is a good starting point, but one needs to verify the predictions using a more detailed model.

We propose a model system of *ABA* polymer translocating through a solid state nanopore under alternating field, where *A* block is a polycation, and *B* is single stranded DNA molecule. Such a proposed set up is intended to serve two main purpose: (a) Once the DNA is pushed into the nanopore by applying high bias, the DNA is expected to move back and forth indefinite number of times under a reduced bias, since the polycation will be prevented from entering the nanopore from either ends (b) at certain alternating frequency of applied voltage, the DNA is expected to ratchet at a slow rate forward steadily. Even if the nucleotides cannot be ratcheted at a steady rate for individual trajectories, useful information may be obtained from the average data.^29,30^ At low frequency of the applied voltage, the nucleotides are expected to drift at a steady rate but faster through the nanopore. ^26^ At very high frequency, the nucleotides are expected to be trapped inside the nanopore.^26,27^ At an intermediate frequency the nucleotides may be ratcheted at a steady forward rate. With these goals in mind, we used a coarse grain model of oxDNA2 model of DNA^32–36^ to investigate the dynamics of nucleotides under an applied alternating voltage. The electric field across the nanopore was determined using Poisson-Nernst-Planck solver and the electric field thus obtained was used later for langevin dynamics simulation of *ABA* translocation through a solid state nanopore.

### Model and Simulation Details

For modeling the translocation of an *ABA* polyelectrolyte through a solid state nanopore, oxDNA2 coarse grain (CG) model^32–36^ was used in LAMMPS.^37,38^ Where single stranded(ss) DNA (*B*) is covalently linked with symmetric polycations (*A*) at both ends. Four types of CG nucleotides are implemented in oxDNA2 to model four different naturally occurring nucleotides, we created two additional types to model the polycation beads and the nanopore beads. The nanopore was made up of immobile excluded volume beads and the polycations were modeled as a bead spring model, each bead had some excluded volume with charge in the center of the bead. The new types are minor modifications of the already existing nucleotide models and the details of these modifications are described in this section. The oxDNA2 model is capable of modeling sequence effects but we focused only on one type of nucleotide (Thymine) in this study. Brief description of the original oxDNA2 model for ssDNA is provided below:

### oxDNA2 Model

This CG model is a nucleotide level model which has been developed in a heuristic manner to reproduce well known properties of ssDNA and double stranded (ds) DNA reported in the experimental literature, such as conformational, thermal and mechanical properties.^36^ The robustness of this model to reproduce experimental results for more complex systems have been demonstrated, for example DNA overstretching and DNA origami.^36^ In oxDNA2 model, each nucleotide is modeled as a backbone bead (representing sugar and phosphate), rigidly bonded and oriented with respect to it’s side group (representing the base). The backbone has one interaction site and the side group has two. The consecutive backbone beads are connected through a finitely extensible non-linear elastic (FENE) potential. Inter-backbone bead interaction includes excluded volume as well as Debye-Hückel interaction potential. Inter-side group bead interactions include stacking, cross-stacking, coaxial stacking, hydrogen bonding and excluded-volume interaction potentials distributed at two different interaction sites. Some of the inter-side group bead interactions happen only when they are non-adjacent, for example hydrogen bonding, cross-stacking and coaxial stacking. The unrelated backbone and side groups interact through excluded-volume potential. The equation form and the parameter value of the potential parameters are provided in Ref.^32,36^ Knowing the potential form and parameter values, net potential of a rigid body can be calculated for a given distribution of the rigid bodies. Each nucleotide is represented as a quaternion containing information about the center of mass and orientation of the nucleotide, which can easily yield the position of the backbone as well as the side group.

Langevin dynamics simulation was used for updating the position and orientation of the quaternions. *n* 3-dimensional rigid bodies can be represented in terms of their center of mass coordinate and rotational coordinates. The center of mass coordinates **r** = (*r*^1*T*^, …, *r*^*nT*^)^*T*^, where 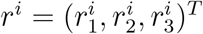 being the cartesian coordinate of the *i*^*th*^ nucleotide and the rotational coordinates in the quaternion representation **q** = (*q*^1*T*^, …., *q*^*nT*^)^*T*^, where 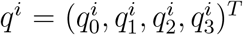 and | *q*^*i*^| = 1, and *T* denotes transpose. Assuming that the potential *U* (**r, q**) is a sufficiently smooth function, such that the net conservative force acting on nucleotide *i* is *f*^*i*^(**r, q**) = −∇_**r**,*i*_*U* (**r, q**) and the net torque due to conservative force on nucleotide *i* is *τ*^*i*^(**r, q**) = Σ _*α*_*d*^*α*^ × (**A**(*q*^*i*^)*f*^*i,α*^), where *d*^*α*^ is the site coordinate of the interaction site *α* relative to the center of mass of nucleotide *i* and **A**(**q**) is the rotational matrix expressed in terms of quaternion coordinates (Note: there is one interaction site corresponding to the backbone and two corresponding to the side group). The net force on a nucleotide *i* is given by:

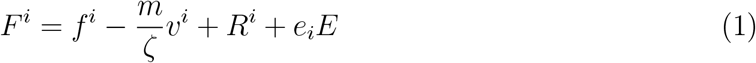

where *f*^*i*^ is the conservative force due to interaction with other nucleotides; second term represents the frictional drag, where *m* is the mass of a nucleotide(same for all nucleotide types), *v*^*i*^ is the velocity and *ζ* is the damping factor; third term represents the force due to random kicks from the implicit solvent and its magnitude is given by 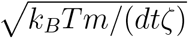 (derived from fluctuation-dissipation theorem) where *k*^*B*^*T* being Boltzmann constant times desired temperature, *dt* is the timestep size; last term represents the electrophoretic driving force, given by the product of the nucleotide backbone charge (*e*_*i*_) times the local electric field. A qualitatively similar form has been presented for the angular momentum, in Davidchack et al,^39^ containing contributions from conservative term due to inter-nucleotide interactions, the frictional and random terms originating from the implicit solvent. An additional parameter, a scaling factor relating rotational to translational friction coefficient is used in the angular momentum calculations. A rigid body langevin-type integrator of the kind “Langevin C” described in Davidchack et al^39^ is used in LAMMPS for updating the position of the center of mass. The quaternion degrees of freedom are updated in LAMMPS through an evolution operator as described in Miller et al.^40^

The original derivation for oxDNA model and parameters has been described in detail in Ref.,^32,34^ the updated model oxDNA2 that has been used in this simulation is described in Ref.^36^ and the details of its implementation in LAMMPS has been described in Ref.^35^

### Modification of oxDNA2 Model

The above scheme works for DNA only system, but the other components in the system such as the polycation tails as well as the nanopore cannot be implemented directly from the model. In order to create a model for the nanopore and the polycations, some modifications were done on the existing nucleotide model. Four types of nucleotides already exists in LAMMPS, two additional types of nucleotides are created: type 5 corresponds to polycation beads and type 6 corresponds to the nanopore beads. In our model, unlike the nucleotides, the polycation beads and the nanopore beads only have one backbone bead and no side group. All the calculations related to the side chain of beads belonging to types 5-6 are turned off. This is achieved by defining stacking potential, hydrogen bonding, cross-stacking and coaxial stacking only between the types 1-4. To turn off the excluded volume interaction between the side groups of type 5 and 6 with others, the distance between the side groups belonging to these types and others were arbitrarily reset to a large value during excluded volume potential calculations. The excluded volume interaction is zero beyond certain cut-off, therefore the excluded volume interaction is zero by default for these pairs. Finally, the Debye-Hückel interaction is defined for types 1-5 and the sign of interaction potential is made negative for pairs: type 5 and types 1-4. Bonds were created for covalently linking the ssDNA and the polycation tails.

For computing the electrophoretic force and torque, the input files and LAMMPS files were modified. In the example file provided for CGDNA model, the charge of the rigid bodies are not defined in the topology file. In the modified version, we defined charges of different types and modified the LAMMPS code to allow reading of charge per rigid body. In LAMMPS, the force term adding the contribution from the electrophoretic term (charge time the local electric field) already exist, and we added codes for calculating torque associated with electrophoresis. The contribution of electrophoresis on the total torque of a rigid body *i* is given by *d*^*bb*^ × (**A**(*q*^*i*^)*e*_*i*_*E*), where *d*^*bb*^ is the site coordinate of the backbone with respect to its center of mass.

### Initial Configuration

The tools provided in LAMMPS USER-CGDNA lets us generate a single stranded helical DNA. We initially generated a ssDNA of type 4 (representing polyT) of certain length. Some part of the generated DNA (polycation tails of predetermined length at both ends) were assigned type 5 to model polycation beads. Next, the immobile nanopore was created by generating center of mass position and quaternion of the nanopore beads (type 6) such that the backbone beads generated from these information leads to a hollow cylinder of predetermined diameter and thickness (Note: There are no side groups associated with the nanopore beads). Besides the hollow cylinder, square walls were created at both ends of the cylinder (excluding the circular areas extended by the cylinder), such that the cylinder and the walls in combination represent a solid state nanopore as shown in Fig. 1. Once such initial configuration was created then the ABA polymer was placed with its center coinciding with the center of the nanopore. Electric field of certain polarity was applied and the ABA polymer was equilibrated under external field. In this new configuration, one end of the DNA lingers near either end of the nanopore, depending on the polarity of the applied voltage. At this stage, one end of the DNA was tethered at the pore end and the chain was equilibrated under no applied electric field. We do not expect a huge difference in qualitative nature of overall polymer conformations of such tethered system with and without applied field because of the limited range of the electric field within and around the narrow nanopore (range ∼ 2 nm). From this stage different initial configuration of the polymer were collected and 600 independent simulations were conducted under an applied field of polarity opposite to that applied during equilibration. According to our convention, the nucleotide indices increases from the right to left, starting at 1 and ending at 20. Initially, one end of the DNA was tethered at *z* = 3 Å such that the DNA does not gets pulled inside the nanopore for the shorter *N*_*A*_. Moreover, the last DNA bead lingers around z coordinate of 0-3 Å at the end of the simulation, hence the end point of a simulation can be simply taken as the start of a new simulation by switching the voltage polarity. The coordinate and the quaternions of the ABA polyelectrolyte were collected at regular time intervals and analyzed.

**Figure 1:**
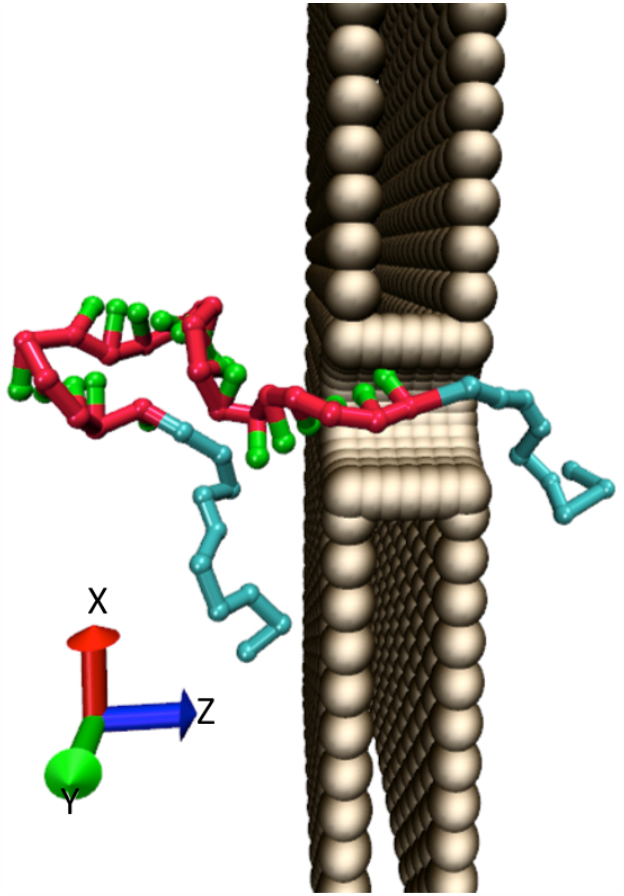
Initial configuration of ABA polyelectrolyte inside a solid state nanopore having one end of the DNA near the right end of the nanopore (initially tethered at *z* =3 Å). Polycation tails (A) are colored cyan, and the polyT (B) has backbone (colored red) and side groups (colored green). The positive electrode is on the right side and the indices of nucleotides increases from right to left according to our convention.

### Poisson Nernst Planck

The voltage (*ϕ*) profile across the nanopore (axis in the z-direction as shown in Fig.1) due to an applied bias in 1M KCl aqueous solution was obtained using Poisson Nernst Planck solver.^41–44^ For an applied bias, the whole simulation box is discretized into grids; Poisson equation and Nernst-Planck equations are numerically solved simultaneously under the constraint of boundary conditions (BCs). The BCs are: the voltage reaches the applied bias on the lateral boundaries, the dissociated ion concentration reaches 1M at the lateral boundaries; periodic boundary conditions are applied in the top/bottom in-plane/out-plane boundaries of the simulation box.

The Poisson equation is given by:

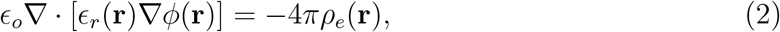

and the Nernst-Planck equation is given by:

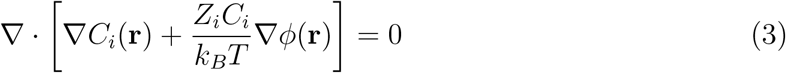

Where *ϵ*_*o*_ is the dielectric permittivity of the free space; *ϵ*_*r*_ is set at 2 for the grids occupied by the nanopore beads and 80 elsewhere; *C*_*i*_(**r**) is the number density of the *i*^*th*^ ion at a location **r**; *i* = *K*^+^, *Cl*^*−*^ and the corresponding *Z*_*i*_ being 1e and -1e, respectively; and the charge density *ρ*_*e*_(**r**) = *e*[*C*_*K*+_(**r**) − *C*_*Cl−*_(**r**)], *e* being electronic charge.

The tolerance for the potential and ion concentrations were set at 10^*−*6^ V and 10^−9^ M, respectively. We used 241×241×181 grids of uniform grid size of 1 Å. The relaxation parameters for the voltage and concentration were 0.004 and 1, respectively.

### Parameters

The nanopore had a diameter of 1.45 nm (after excluding the volume occupied by the pore beads) and a thickness of 1.53 nm. Two different kinds of voltage were applied across the nanopore: direct and alternating. The amplitude of the voltage ranged from 0.18 V to 0.72 V and for the alternating (square wave type) voltage, the time period of oscillation was varied around the most probable residence time of a nucleotide (determined from the direct voltage simulations). The number of nucleotides in the ssDNA (B block) was set at 20 and the number of beads in the A block were: 2, 10, 30 and 50. In total 600 simulations were conducted per case for statistical analysis.

The dynamics of the nucleotides were investigated in two different ways : (i) mean displacement of nucleotide *i* along z-axis ⟨*z*⟩ as a function of time (ii) most probable time of finding nucleotide *i* within -1.5 Å - 1.5 Å (∆*z*). By position of the nucleotide *i*, we mean the position of its backbone. For every simulation, the first passage time of the *i*^*th*^ nucleotide into the left side of the nanopore (−7.65 Å) (*t*_*o*_) was determined (Note: due to simulation constraints we start with some nucleotides being already within the nanopore; they were not considered for the mean displacement calculations). At this stage the z-displacement of the *i*^*th*^ backbone was monitored as a function of *t* − *t*_*o*_. The mean z-displacement was obtained by averaging over 600 independent simulations. For determining the most probable time of finding the *i*^*th*^ nucleotide within ∆*z*, individual simulations were considered first and the probability of finding *i*^*th*^ nucleotide within ∆*z* was determined (either 1 or 0) as a function of time. 600 independent probability distributions were averaged to obtain the mean probability distribution (*p*_*i*_(∆*z*)) as a function of time, which is then gaussian fitted to obtain the peak time (most probable time) for appearance of *i*^*th*^ nucleotide within ∆*z*.

## Results and Discussion

The goal of this simulation is to determine how accurately we can monitor the nucleotide motion along the nanopore by electrophoresis of ABA polymer through a solid state nanopore, where B is the ssDNA (polyT) and A is a polycation. With this setup, the capture of ABA can be initiated at high bias, the bias can then be reduced once the B block is fully within the nanopore. The capture of ABA polymer by the nanopore and the entry of B into the nanopore is beyond the scope of this study, here we investigated the dynamics of the nucleotides once B is placed within the nanopore. The electrostatic potential corresponding to applied voltage across the nanopore is determined by using Poisson-Nernst-Planck solver. Cubic spline interpolation was used to get a potential profile on a finer grid and then numerical derivative with respect to distance was done to obtain electric field profile across the nanopore as shown in Fig. 2. The electric field as a function of distance was converted to LJ units and then fitted with polynomial equations. Finally, the fitted polynomial equations were used as an input parameter for the langevin dynamics simulation calculations in LAMMPS. The electric field within the nanopore was in the range of 2-12 mV/Å, it sharply dropped right next to the nanopore boundary and then slowly approached zero on both sides of the nanopore after a distance of about 2 nm from the nanopore boundary. Note that the electric field reported here is half of the actual value because the effective charge per nucleotide in the model is 0.815*e* (where *e* is the electronic charge) for Debye-Hückel interaction among the charged species. The magnitude of charge used in the model is optimized with other interaction terms in the model, but the electrophoretic force experienced by the nucleotides is much smaller due to smaller effective charge ∼ 0.4,^45^ hence the electric field is halved to reduce the electrophoretic force without interfering with the default interaction potentials as prescribed by the oxDNA2 model.

**Figure 2:**
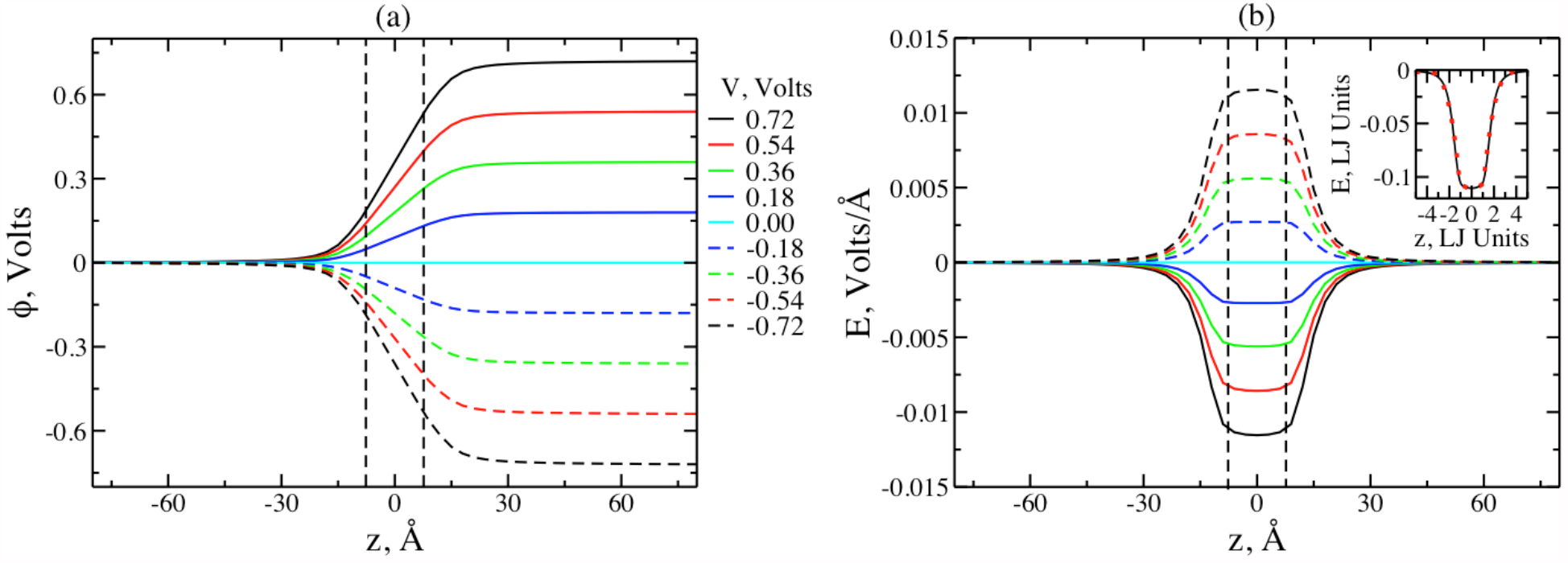
(a) Electrostatic potential (*ϕ*) across the simulation box corresponding to different applied voltage (*V*). (b) Electric field (*E*) across the simulation box for different *V*. The inset shows electric field profile (in LJ units) as a function of the z-coordinate (in LJ units) fitted to polynomial functions (red circles) corresponding to applied voltage of 0.54 volts. Cubic polynomial fitting was done within the nanopore and fourth order polynomial fits were done on left and right side of the nanopore. The fitted polynomial functions of *E* (LJ units) as a function of *z* (LJ units) were taken as an input for the langevin dynamics simulations. The two vertical dashed black lines show the boundary of the nanopore in both the figures.

### Influence of Direct Voltage on Nucleotide Dynamics

We investigated the effect of *N*_*A*_ on the displacement of individual nucleotides within the nanopore. As shown in Fig. 3 (a)-(d), the nucleotide displacement is linear for most part of the DNA (till 16^*th*^ nucleotide); the 20^*th*^ nucleotide exits the pore very quickly for *N*_*A*_ =2 but it rattles about *z* = 0-4 Å for higher values of *N*_*A*_. The average residence time within the nanopore is between 0.01 - 0.015 *µ*s for *i* ≤ 16. We also investigated the effect of *V* on the displacement of nucleotides as shown in Fig. 3 (e)-(h). The average displacement was linear function of time for *i* ≤ 16 and the residence time of the nucleotides decreased with increase in *V*. The average residence time decreased sharply for lower *V* (0.03 *µ*s to 0.01 *µ*s corresponding to *V* = 0.18 and 0.36 volts, respectively) and saturated for higher *V* (about 0.006 *µ*s for *V* = 0.54 and 0.72 volts). The 20^*th*^ nucleotide lingered around 0 Å and 3 Å for *V* = 0.18 volts and 0.36/0.54 volts, respectively; it wildly fluctuated between 0-7 Å for *V* = 0.72 volts. Such wild fluctuations may facilitate DNA escape from the nanopore. For a very similar nanopore (3nm thick and 2 nm diameter) and for an applied voltage of 300 mV (corresponding to about *V* =0.54 volts in our case), Ref.^26^ reported a residence time/base of 0.49 ns based on all atomistic MD simulations, which is at least an order of magnitude smaller than our result. The average residence time per nucleotide from our calculations varied between 6 - 30 ns. As our pore diameter is slightly smaller than theirs (1.45 nm versus 2 nm^26^) this can make a huge impact on the residence time. The residence time for a ssDNA in a 20 nm thick nanopore, calculated using MD simulations, increased by an order of magnitude from 0.18 to 1.4 *µ*s/base by decreasing the pore diameter from 2.3 to 1.4 nm. ^46^ For a very similar sized nanopore (1.4-2.1 nm diameter and 0.5-8.9 nm thickness) and a high *V* = 0.9 volts, Ref.^16^ reported a residence time of 2-21 ns/nucleotide using ionic current based nanopore experiments, agreeing with our simulation results.

**Figure 3:**
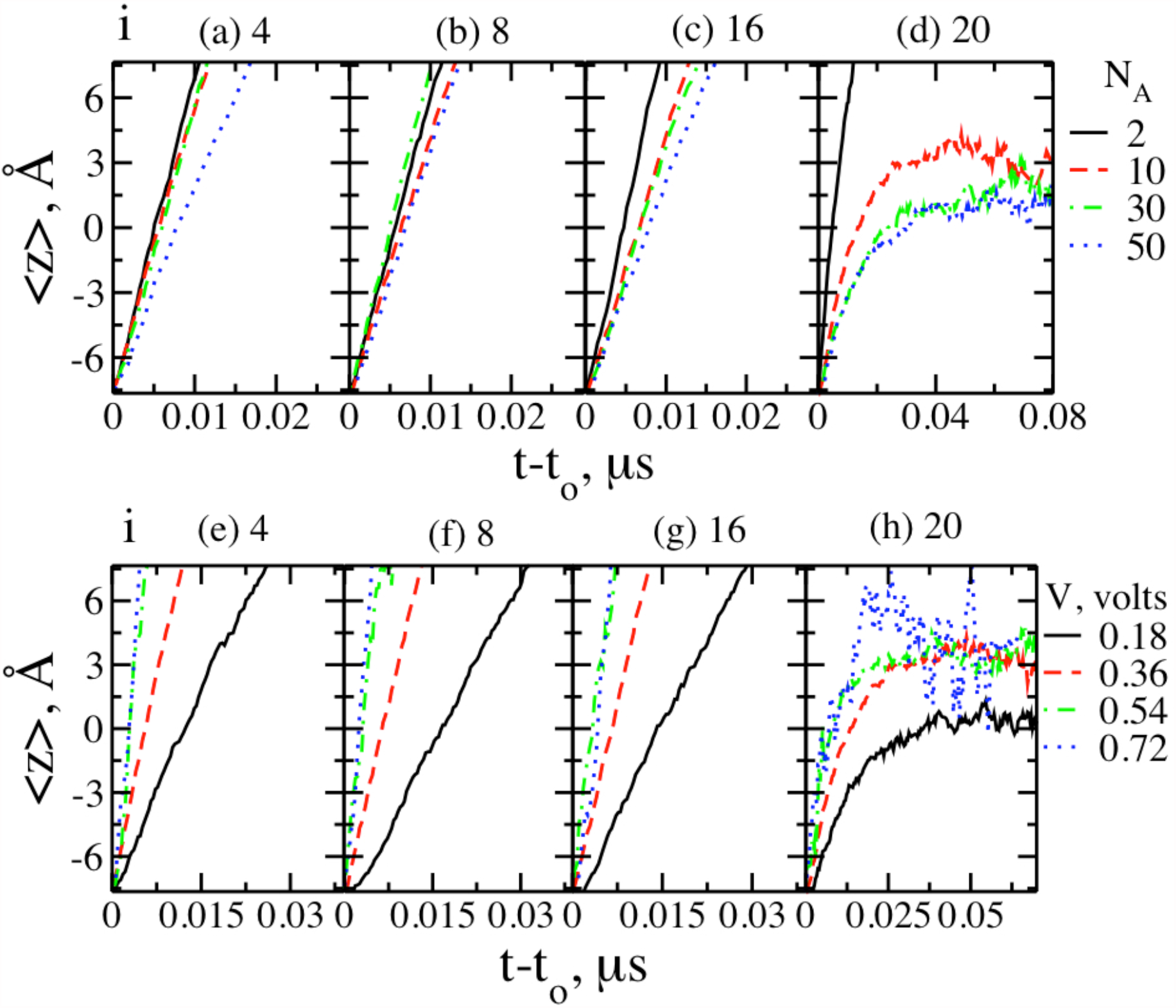
The effect of two different parameters on the dynamics of nucleotides inside the nanopore extending from -7.65 Å to 7.65 Å ; the nucleotide indices (*i*) are shown on top of the corresponding figures. (a)-(d) shows dynamics of different nucleotides as a function of *N*_*A*_ at *V* =0.36 volts. (e)-(h) shows the dynamics of those same nucleotides as a function of *V* corresponding to *N*_*A*_=10. For (a)-(c) the nucleotide displacement are linear and the average residence time is between 0.01-0.015 *µ*s but for (d), the nucleotide escaped for very short *N*_*A*_ =2; it rattled between 0-4 Å for larger values of *N*_*A*_. For (e)-(g), the nucleotide displacements are linear and the average residence time decreased from approximately 0.03 *µ*s to about one quarter of its value with increase in *V* from 0.18 to 0.72 volts. It increased sharply for smaller *V* and saturated for higher values of *V*. (h) showed a nonlinear behavior for all the cases, nucleotides ratcheted around 0 Å for *V* = 0.18 volts and approximately 3 Å for *V* = 0.36/0.54 volts and it wildly fluctuated between 0-7 Å for *V* =0.72 volts, showing a tendency to escape the nanopore at higher applied voltage.

The purpose of the displacement analysis was to track the motion of nucleotides once they are within the nanopore, but it does not give any information about the time difference between the appearance of consecutive nucleotides at the nanopore. Therefore, we analyzed our simulations in a different manner, by tracking the probability of appearance of a nucleotide *i* at any given time in the center of the nanopore ∆*z* (between -1.5 Å and 1.5 Å). Once the 600 independent probability distribution functions were obtained, mean probability distributions were obtained in smaller batches first. A batch of 100 simulations was enough to create a gaussian distribution similar to that shown in Fig. 4(a). Here we present a smoother mean probability distribution plot based on 600 independent probability distributions. As we progress along the DNA length, the probability distribution became wider due to diffusion. Gaussian fitting was performed on the probability distribution to obtain the most probable time of appearance of nucleotide *i* in ∆*z*. The peak times versus the nucleotide index (*i*) showed a linear dependence for a range of applied voltage from 0.18 to 0.72 volts (Fig. 4(b)). Such linear dependence did not last throughout the length of the DNA, but the electrophoretic pull of the polycation slowed down the drift velocity of the nucleotides for larger *i* values (not shown). Moreover, enough data was not available for creating a smooth probability distribution curve for very large *i* values. We obtained the most probable residence time of the nucleotides by determining the slope of the linear fits as shown in inset of Fig. 4(b). The residence time, *τ* decreased sharply for lower *V* values and saturated at higher *V*, which is consistent with our displacement analysis results. But *τ* values are 2-3 times smaller than that obtained using displacement analysis due to a long tail observed in the probability distribution. The purpose of gaussian fitting in this analysis was just to obtain the peak position, hence the contribution from the tail does not appear in this analysis unlike the average displacement.

**Figure 4:**
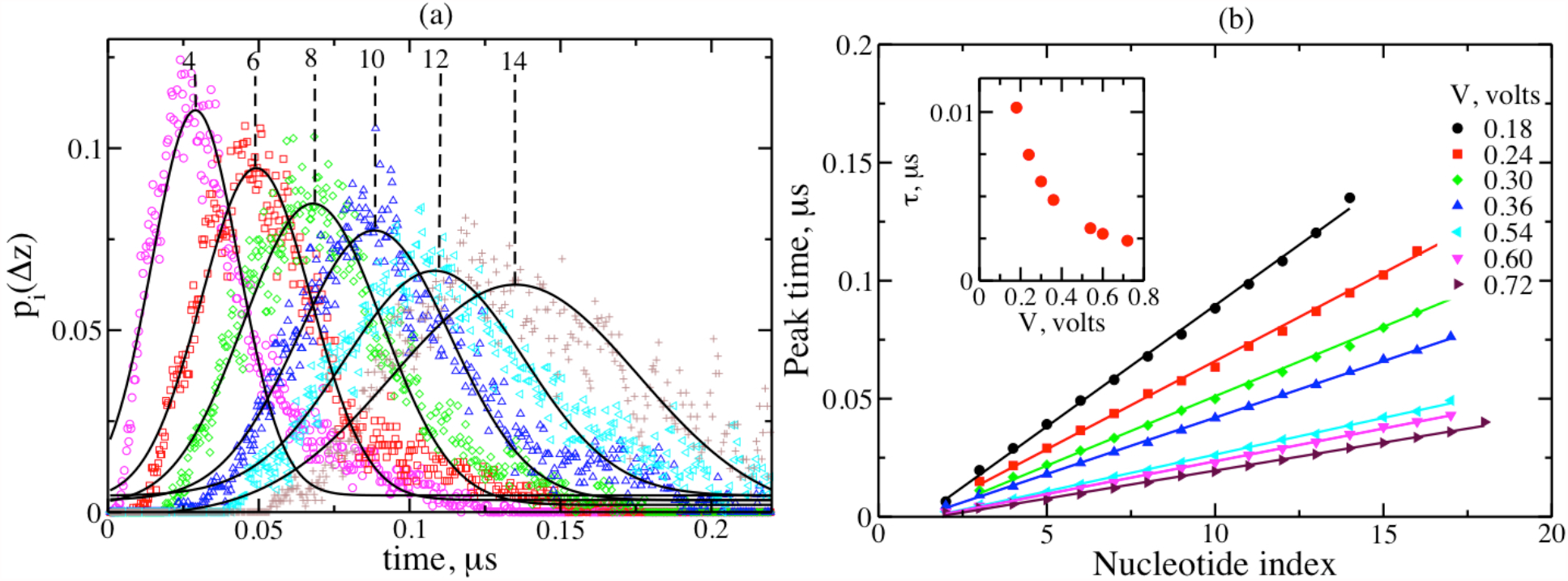
(a) Mean probability distribution (*p*_*i*_) of finding a nucleotide *i* within ∆*z* (−1.5 Å -1.5Å along the z-axis) for *N*_*A*_ = 10 and *V* = 0.18 volts; all simulations were initiated by tethering *i* =1 at *z* =3 Å. The mean probability distribution was obtained by averaging over 600 independent probability distributions and gaussian fitted (solid curves) to determine the peak time. (b) The peak time (determined from (a)) as a function of nucleotide indices for different voltages (*N*_*A*_ = 10 for all the cases) shows linear dependence on *i* for *i* = 2-15 for most cases and upto 18 for higher *V*. The slope of the lines can be used to estimate most probable residence time per nucleotide (*τ*). The inset in (b) shows *τ* as a function of *V* which decreases from 0.0102 *µs* to about 0.0024 *µs* with increase in *V*, showing sharper drop for smaller *V* and saturating at higher *V*.

These calculations showed that the DNA can be retained inside the nanopore by attaching polycations at its ends with *N*_*A*_ ≥ 10 and for *V <* 0.72 volts. Secondly, this configuration enables repeated analysis of the trapped DNA. With repeated analysis, the most probable time of appearance of different nucleotides within the nanopore can be obtained, which shows linear dependence on the nucleotide indices. It should be noted that *p*_*i*_(∆*z*) was obtained after gaussian fitting, the real simulation data is not as smooth, resulting in significant overlap between consecutive peaks at higher *i*. The diffusive noise makes the precise monitoring of the nucleotides using such analysis difficult. Moreover, the average residence time was also too small to be of practical use (*τ* ≤ 0.01 *µs*). As proposed by Mondal and Muthukumar,^31^ the signal to noise ratio can be improved by applying alternating external force, we performed simulation with alternating voltage and applied similar analysis.

### Influence of Alternating Voltage on Nucleotide Dynamics

We investigated the effect of alternating voltage frequency and amplitude on the dynamics of nucleotide, keeping *N*_*A*_ = 10 fixed for all the cases. The dimensionless frequency (*f*) is the ratio of the residence time (*τ*) of a nucleotide over the time period of oscillation. The residence time *τ* was obtained from Fig. 4(b); square wave type of signal was used for alternating voltage, where the amplitude varied between +*V* and -*V* at regular time period. Fig. 5 (a)-(d) shows the average displacement of different nucleotides (*i* value shown at the top of corresponding figures) as a function of time corresponding to different *f* for a fixed *V* = 0.36 volts. The displacement of the nucleotides between the dashed lines (nanopore boundaries) for the 4^*th*^ and 8^*th*^ nucleotide show qualitatively similar behavior (12^*th*^ nucleotide also showed similar behavior, not shown for clarity) but differ from 16^*th*^ and 20^*th*^ nucleotides. For *i* ≤ 12, the qualitative nature of the displacement changed from linear to undulations at regular intervals as *f* was increased from 0.27 to 0.65. With further increase in *f*, the magnitude and frequency of the undulations became smaller (appearing almost linear) and at the highest *f* = 4.8, the displacement was observed to be of unpredictable nature. Relative to the direct voltage or alternating voltage with low *f* = 0.27, the average residence time of the nucleotides inside the nanopore increased by an order of magnitude from ∼ 0.01 *µs* to ∼ 0.1 *µs* as *f* increased. Corresponding to *i* = 16, the nucleotides escaped for *f* = 0.27 and 0.65, but they rattled inside the nanopore for *f* = 1.3, and for higher values of *f* they appear to be rattling inside the nanopore initially and getting out through the nanopore entrance at larger times. For *i* =20, the nucleotides corresponding to *f* = 0.27 and 0.65 rattled within the nanopore and for higher frequencies they bounced away from the nanopore entrance.

**Figure 5:**
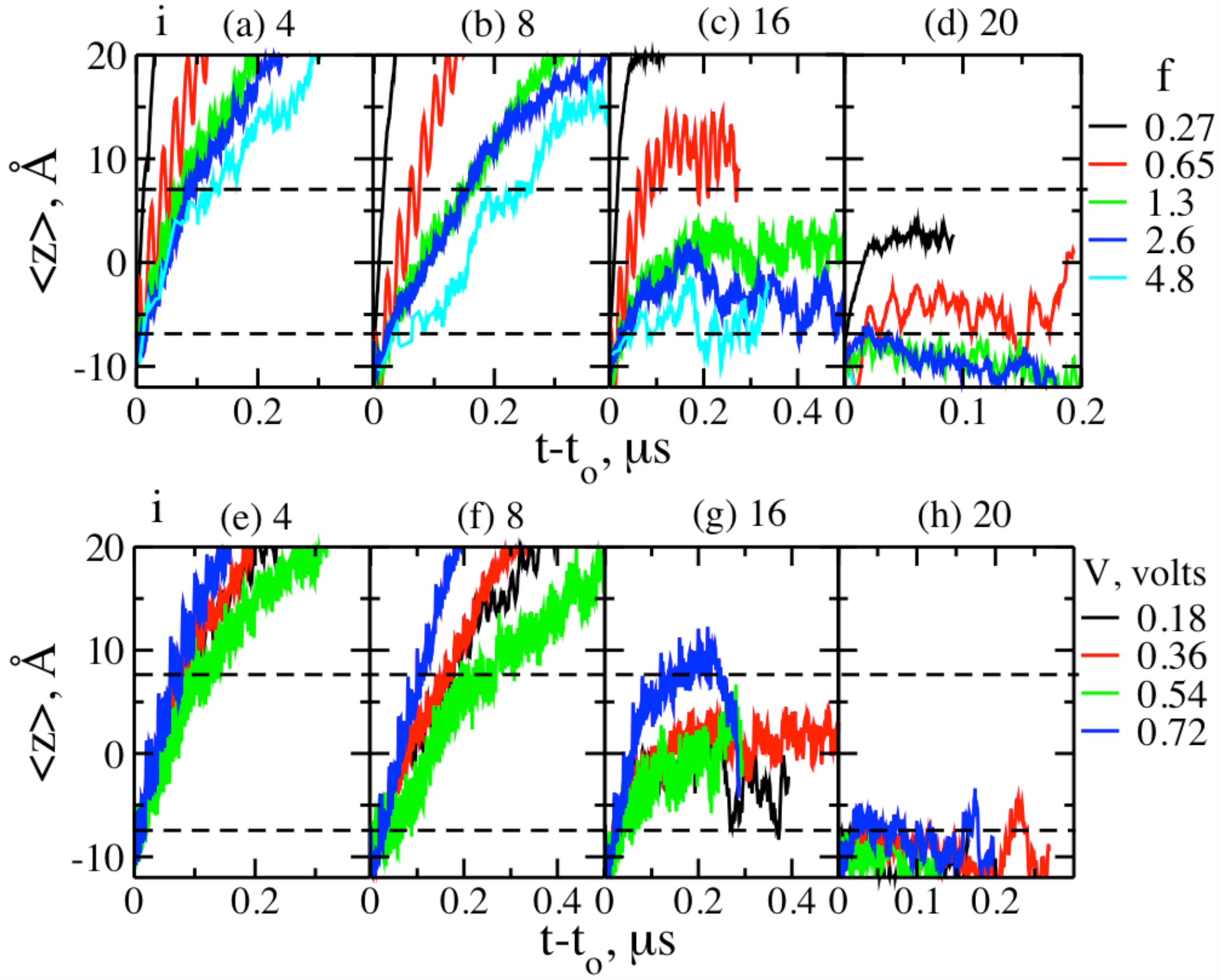
(a)-(d) shows nucleotide dynamics as a function of applied alternating voltage frequency for *V* = 0.36 volts. The qualitative nature of the displacement changed from linear to undulated as the frequency of the applied voltage was increased for smaller *i* (values showed on top of corresponding figures) and the residence time within the nanopore increased by an order of magnitude as well. For larger *i*, they either exited, rattled within or bounced back from nanopore. (e)-(h) shows effect of voltage for constant *f* =1.3. The residence time for the smaller nucleotide indices (4 and 8) showed non-monotonic behavior, no change for 0.18-0.36 volts, increased from 0.36 -0.54 volts, decreased from 0.54-0.72 volts. The 16^*th*^ nucleotide either ratcheted for lower *V* or bounced back from nanopore exit corresponding to *V* = 0.72 volts. The last nucleotide bounced back from the nanopore entrance for all the cases. The two dashed horizontal lines indicate the boundary of the nanopore.

The effect of voltage amplitude on the nucleotide displacement is presented in Fig. 5(e)-(h) for different nucleotides (values displayed on top of the figure) for a fixed *f* = 1.3. As mentioned in the previous case, the 12^*th*^ nucleotide is not shown for clarity but it behaved the same as the 8^*th*^ nucleotide. For *i* ≤ 12, the nucleotide displacement showed a non-monotonic dependence on *V*. The average displacement versus time showed no dependence on *V* at lower *V* = 0.18/0.36 volts, residence time increased for intermediate *V* = 0.54 volts and residence time decreased for higher *V* = 0.72 volts. For *i* = 16, the nucleotides rattled inside the nanopore for all the cases except for the highest voltage. For *V* = 0.72 volts, the nucleotide lingered at the nanopore exit for about 0.1 *µ*s and bounced back into the nanopore afterwards. For *i* = 20, all the nucleotides bounced back from the nanopore entrance. The effect of alternating voltage on the nucleotide dynamics have been reported in literature.^26,27^ Corresponding to the pore dimension similar to ours,^26^ the middle nucleotide rattled inside the nanopore for a very high frequency of 2500 MHz and 300 mV. Similarly, Sigalov and coworkers^27^ also demonstrated rattling motion in a longer (10.3 nm thick) hour- glass nanopore corresponding to a variable frequency (≥ 500 MHz) and 20 volts amplitude. In our simulations, for the case of *f* = 0.65 and *V* = 0.36 volts correspond to 125 MHz frequency and our nanopore is an order of magnitude smaller than Ref.,^27^ we observed rattling of nucleotides for higher *i* at higher *f*. When it reached the end of the DNA (higher *i*), the polycation tail influenced the nucleotides to rattle inside the nanopore, irrespective of *f*.

The average displacement of nucleotides showed encouraging results in terms of enhanced residence time/nucleotide (of the order of 0.1 *µs*, still low for experimental detection) and almost linear displacement as a function of time at intermediate *f* = 1.3/2.6 for *i* ≤ 12. But only the gaussian fit shows encouraging results, the real data is noisy and the peak time of the consecutive nucleotides overlap significantly. To understand why even after averaging over 600 independent simulation we cannot obtain distinct simulation peaks for the nucleotides, we examined the individual trajectories. The individual trajectories of the nucleotides show stochastic behavior for all the cases as shown in Fig. 6. Fig. 6(a)-(d) shows effect of *f* for a fixed *V* = 0.36 volts and Fig. 6(e)-(h) shows effect of *V* for a fixed *f* = 1.3. At low frequency (*f* = 0.27, in absolute units it is about 50 MHz), the nucleotide motion is stochastic and its average residence time is low (∼ 0.01 *µs*). Using a similar sized nanopore, Ref.^16^ also reported variability in the dwell time for 10 MHz frequency. At very high frequency, (2500 MHz^26^ and ≥ 500 MHz^27^) any nucleotide can be trapped within a nanopore for a long time. At intermediate frequency (∼ 125 MHz), the average residence time of nucleotides can be enhanced by an order of magnitude to ∼ 0.1 *µs*, but the system became more stochastic. The longer a nucleotide resides within a nanopore, the chain conformation outside the nanopore can evolve into different states. Some conformations may favor the forward motion but others may not, hence such variability in the displacement behavior of individual trajectories is observed.

**Figure 6:**
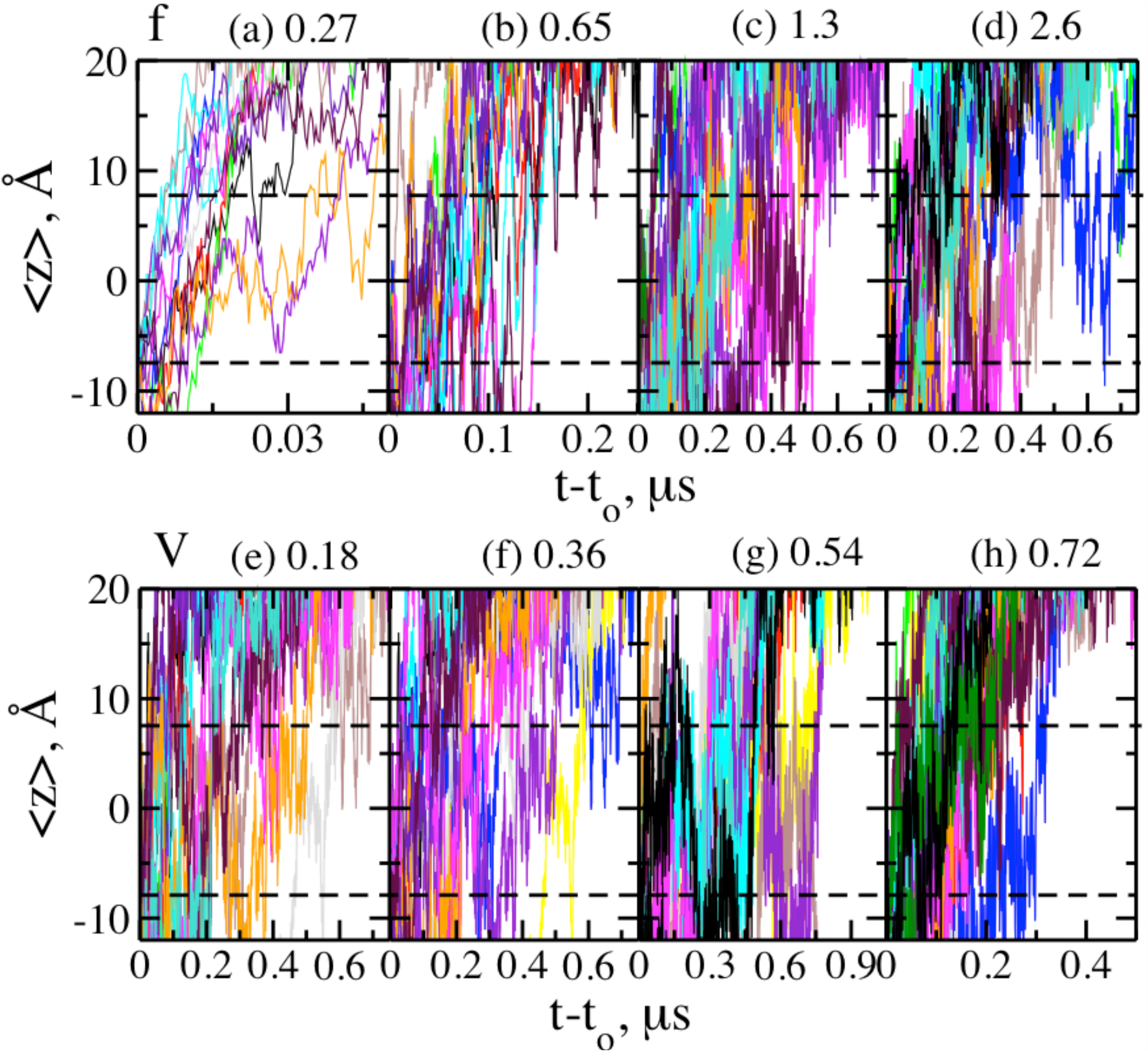
Individual trajectories of the displacement of the 8^*th*^ nucleotide as a function of time within the nanopore. (a)-(d) shows the effect of *f* for *V* = 0.36 volts and (e)-(h) shows effect of *V* for *f* =1.3. The individual trajectories are stochastic, less so for lower frequency which corresponds to lower residence time.

Since such design easily allows for repeated analysis, can we get some meaningful information from repeated analysis instead of futile attempt on trying to analyze individual trajectories? Repeated analysis has been successfully applied on the dsDNA as well as ssDNA translocation through a double nanopore setup. ^29,30^ We repeated the probability analysis as described in the direct voltage section. In this technique, we monitored the most probable time of appearance of a nucleotide *i* within ∆*z* (between -1.5 Å and 1.5Å). With increase in *f*, the probability distribution became wider (not shown) and the peak time versus nucleotide indices are not so linear compared to lower *f* as shown in Fig. 7(a). Fig. 7(a) shows that the peak time versus nucleotide index fits well to a linear regression for lower *f* = 0.27, for higher *f* they are still linear but the fluctuations are larger. Moreover, the number of nucleotides that follows this approximate linear behavior also reduced with increase in *f*. Lastly, there were insufficient number of data points for gaussian fitting corresponding to higher *i* and higher *f*. As shown in the inset Fig. 7(a), the slope of the lines (measure of most probable residence time *τ*) increased sharply from *f* = 0.27 till *f* = 1.3, *τ* did not increase as sharply when *f* was further increased to 2.6. Similar to the direct voltage case, the most probable residence time is lower than the average residence time by a factor of 2-3, estimated from Fig. 5(a) due to longer tail in the mean probability distribution.

**Figure 7:**
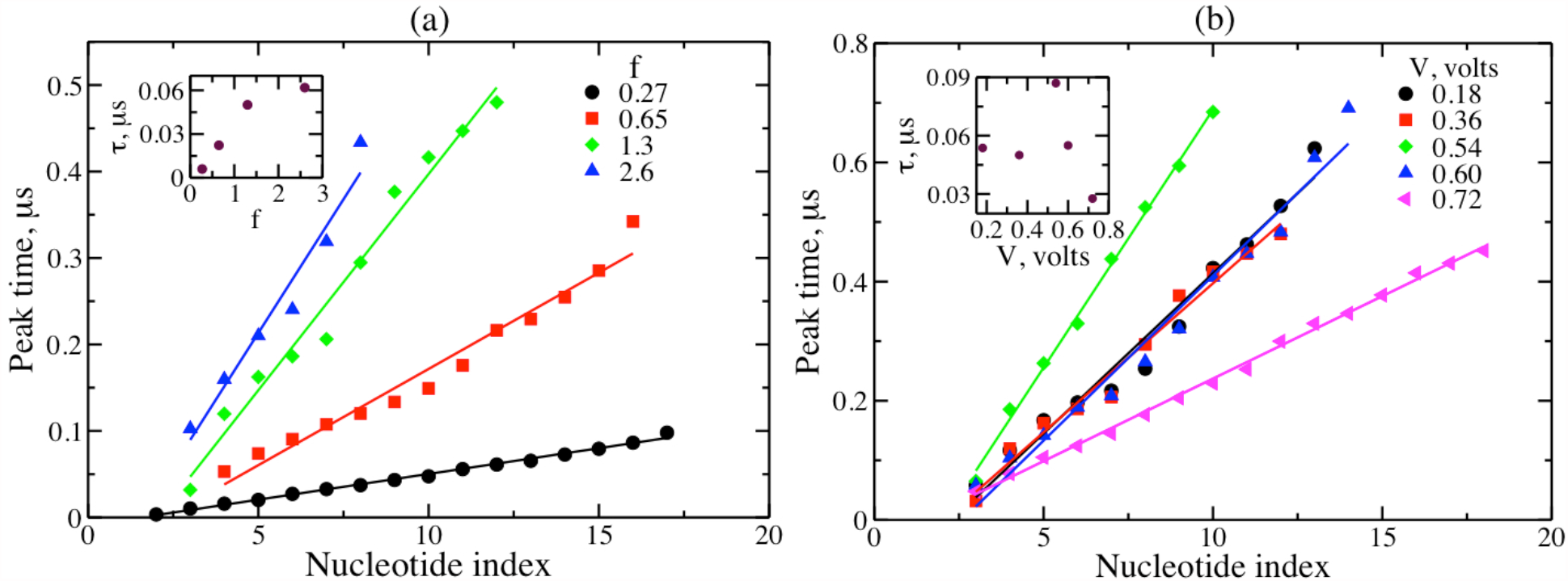
(a) Effect of *f* on the peak time obtained by gaussian fitting scheme described in Fig. 4(a) corresponding to *V* =0.36volts. The peak time is linear for lower frequency, and is fluctuating for higher frequencies. The residence time increased sharply for lower values of *f* and saturated at higher *f*. (b) Effect of *V* on the peak time for *f* =1.3 showing a non-monotonic behavior. The residence time almost remained the same for *V* =0.18 and 0.36 volts, increased at *V* =0.54 volts and decreased with further increase in *V*.

We also investigated the effect of *V* for a fixed *f* = 1.3 as shown in Fig. 7(b). The most probable residence time (*τ*) showed non-monotonic behavior: did not change for *V* = 0.18 and 0.36 volts, increased from 0.05 to 0.09 *µs* for increase in *V* from 0.36 volts to 0.54 volts, decreased to 0.028 *µs* for *V* = 0.72 volts. For understanding such a non-monotonic behavior we followed the electric field profile as a function of voltage (Fig. 2(b)). At high *V*, the nucleotides within the nanopore as well as in the vicinity of the nanopore experiences strong field, hence some of the nucleotides are expected to be in stretched conformation at least in the vicinity of the nanopore, such favorable conformation in the vicinity of the nanopore may facilitate relatively faster translocation on an average. At low *V*, the nucleotides within the nanopore experiences very small field and the nucleotides in the vicinity of the nanopore experience very weak field. The nucleotides outside the nanopore are able to explore all possible conformations. At intermediate *V*, the nucleotide right next to the nanopore experiences a field that facilitates its orientation along the nanopore but the rest of the nucleotides are able to explore all possible conformations due to sudden drop in field. Such discord right at the nanopore boundary may create unfavorable conformations leading to slow down of translocation on an average.

In summary, we explored the possibility of using ABA polyelectrolyte translocation under alternating voltage as a possibility for genome sequencing. Using alternating voltage at high frequency, we were able to increase the average dwell time to ∼ 0.1 *µs* (an order of magnitude higher than the direct voltage) but the highly stochastic nature of the polyelectrolyte dynamics does not allow us to monitor the position of the nucleotides precisely.

## Conclusions

A novel strategy for genome sequencing has been designed. In this strategy, the negatively charged single stranded DNA (*B*) is covalently bonded with positively charged (*A*) symmetric polymers at both the ends. When such chain is captured inside a nanopore, with DNA block inside the nanopore, the ssDNA can be electrophoretically driven back and forth. The effect of external voltage amplitude, voltage frequency and tail length were investigated on the dynamics of the nucleotides. We found that for a polycation length of 10 or higher and a voltage lower than 0.72 volts, the DNA can be retained within the nanopore and driven back and forth indefinite number of times. The average residence time of a nucleotide within the nanopore was too small to be of any practical use (6-30 ns) and the stochastic nature of the polymer dynamics made it difficult to monitor the position of the nucleotides within the nanopore, precisely. As proposed by Mondal and Muthukumar,^31^ alternating eternal force may allow for precise monitoring of the nucleotides due to stochastic resonance, hence we used square wave type alternating voltage for precise control of nucleotide dynamics. Using alternating voltage, we increased the average residence time of nucleotides by an order of magnitude but the individual trajectories were stochastic. Since the DNA can be repeatedly analyzed easily, therefore we proposed to construct most probable time of appearance of nucleotides with the nanopore. With such construct we were able to get almost linear dependence of most probable time versus nucleotide index only through gaussian fitting.

## Acknowledgement

Acknowledgement is made to the National Institutes of Health (Grant No. 5R01HG002776- 16), the National Science Foundation (DMR-2015935), and the AFOSR Grant FA9550-20- 1-0142 for financial support.

